# *Candida glabrata* maintains two Hap1 homologs, Zcf27 and Zcf4, for distinct roles in ergosterol gene regulation to mediate sterol homeostasis under azole and hypoxic conditions

**DOI:** 10.1101/2024.06.20.599910

**Authors:** Debasmita Saha, Justin B. Gregor, Smriti Hoda, Katharine E. Eastman, Mindy Navarrete, Jennifer H. Wisecaver, Scott D. Briggs

## Abstract

*Candida glabrata* exhibits innate resistance to azole antifungal drugs but also has the propensity to rapidly develop clinical drug resistance. Azole drugs, which target Erg11, is one of the three major classes of antifungals used to treat *Candida* infections. Despite their widespread use, the mechanism controlling azole-induced *ERG* gene expression and drug resistance in *C. glabrata* has primarily revolved around Upc2 and/or Pdr1. In this study, we determined the function of two zinc cluster transcription factors, Zcf27 and Zcf4, as direct but distinct regulators of *ERG* genes. Our phylogenetic analysis revealed *C. glabrata* Zcf27 and Zcf4 as the closest homologs to *Saccharomyces cerevisiae* Hap1. Hap1 is a known zinc cluster transcription factor in *S. cerevisiae* in controlling *ERG* gene expression under aerobic and hypoxic conditions. Interestingly, when we deleted *HAP1* or *ZCF27* in either *S. cerevisiae* or *C. glabrata,* respectively, both deletion strains showed altered susceptibility to azole drugs, whereas the strain deleted for *ZCF4* did not exhibit azole susceptibility. We also determined that the increased azole susceptibility in a *zcf27Δ* strain is attributed to decreased azole-induced expression of *ERG* genes, resulting in decreased levels of total ergosterol. Surprisingly, Zcf4 protein expression is barely detected under aerobic conditions but is specifically induced under hypoxic conditions. However, under hypoxic conditions, Zcf4 but not Zcf27 was directly required for the repression of *ERG* genes. This study provides the first demonstration that Zcf27 and Zcf4 have evolved to serve distinct roles allowing *C. glabrata* to adapt to specific host and environmental conditions.

**IMPORTANCE:** Invasive and drug-resistant fungal infections pose a significant public health concern. *Candida glabrata*, a human fungal pathogen, is often difficult to treat due to its intrinsic resistance to azole antifungal drugs and its capacity to develop clinical drug resistance. Therefore, understanding the pathways that facilitate fungal growth and environmental adaptation may lead to novel drug targets and/or more efficacious antifungal therapies. While the mechanisms of azole resistance in *Candida* species have been extensively studied, the roles of zinc cluster transcription factors, such as Zcf27 and Zcf4, in *C. glabrata* have remained largely unexplored until now. Our research shows that these factors play distinct yet crucial roles in regulating ergosterol homeostasis under azole drug treatment and oxygen-limiting growth conditions. These findings offer new insights into how this pathogen adapts to different environmental conditions and enhances our understanding of factors that alter drug susceptibility and/or resistance.

## INTRODUCTION

Invasive and drug resistant fungal infections are significant public health issues and new estimates indicate that life-threatening fungal infections affect over 6.5 million people globally each year (1). Among these global invasive fungal infections, more than 70% are caused by invasive *Candida* species which include *Candida albicans* and other non-*albicans* (NAC) *Candida* species such as *C. glabrata*, *C. krusei*, *C. tropicalis*, and *C. parapsilosis* (2–5). Of the NAC species listed, *Candida glabrata* is considered the second or third most commonly isolated NAC *Candid*a species, with *C. albicans* being the most commonly isolated (2, 4–6). The traditional genus *Candida* is a paraphyletic group, and *C. glabrata* is more closely related to *S. cerevisiae* than to other common human pathogens including *C. albicans* (7). The last common ancestor (LCA) of *C. glabrata* and *C. albicans* existed ∼250 million year ago (Mya) whereas the LCA of *C. glabrata* and *S. cerevisiae* occurred ∼50 Mya (8). *C. glabrata* is considered the major pathogenic species of the post-whole genome duplication (WGD) *Saccharomycetaceae* group, with immunosuppressed patients (e.g., those with diabetes mellitus, cancer, or organ transplants) and/or elderly patients being particularly susceptible to these infections (6, 9–12).

*C. glabrata* (*Cg*) is also a non-CTG clade *Candida* species that is known for its intrinsic resistance to azole drugs and ability to develop clinical azole drug resistance (7, 13, 14). Azole drugs target and inhibit the enzyme lanosterol 14-α-demethylase (Erg11) which is an essential enzyme for the production of ergosterol in fungi (15–17). Mechanisms of acquiring clinical azole drug resistance have been extensively documented across *Candida* species and include mutations in *ERG11*, *ERG3*, *UPC2* and/or *PDR1* (14, 18–25). Among these genes, gain of function (GOF) mutations in the zinc cluster transcription factors Upc2 and Pdr1 result in increased expression of *ERG11* and/or the ABC drug transporter *CDR1,* respectively (24, 26–30). As for *C. glabrata* clinical drug resistant isolates, Pdr1 GOF mutations are considered the predominant cause for clinical drug resistance (28, 31).

In addition to Upc2 and Pdr1, several known and/or putative zinc cluster factors (Zcf) are critical transcriptional regulators involved in stress response in fungi and amoeba (32, 33). Interestingly, 17 of the 41 *C. glabrata ZCF* genes when deleted show enhanced azole susceptibility as indicated by MIC and/or plate-based growth assays (34). This observation underscores the importance and need to further investigate the role of these zinc cluster transcription factors in *C. glabrata*. However, with the exception of Upc2A (*CgZCF5*), Pdr1 (*CgZCF1*), Stb5 (*CgZCF24*), and Mar1 (*CgZCF4*), little research has been done to understand the mechanistic role of other *C. glabrata* Zcf proteins during azole treatment conditions and/or hypoxic growth (30, 35–39).

In this report, we show for the first time that *S. cerevisiae* (*Sc*) strains deleted for *HAP1* exhibit azole hypersusceptibility when compared to a FY2609 WT strain containing a WT copy of the *HAP1* gene. Interestingly, S288C strains, including the commonly used BY4741 and BY4742, exhibit similar azole susceptibility to *hap1Δ* strains due to a partially disrupted *hap1* gene by a *Ty1* element (*hap1-Ty1* mutant). Based on these observations, we hypothesized that deletion of *C. glabrata HAP1* homologs would also have a similar azole susceptible phenotype. Our phylogenetic analysis indicated that *C. glabrata* contains two proteins, Zcf27 and Zcf4, which are homologs of *S. cerevisiae* Hap1. However, only deletion of *C. glabrata ZCF27,* but not *ZCF4,* showed an azole hypersusceptible phenotype. Upon further investigation, we established that altered azole susceptibility of the *zcf27Δ* strain is attributed to a decrease in azole-induced *ERG* gene expression, resulting in a subsequent reduction in total ergosterol levels. Moreover, azole hypersusceptibility of the *zcf27Δ* strain was alleviated when complemented with a plasmid expressing *ZCF27* or when exogenous ergosterol was introduced into the growth media, but not when the *AUS1* sterol transporter was deleted. Interestingly, unlike Zcf27, Zcf4 protein was nearly undetectable under both untreated and azole-treated conditions. However, under hypoxic conditions Zcf4 was highly induced, while the expression of Zcf27 remained unchanged.

Moreover, the *zcf4Δ* strain showed a growth defect under hypoxic conditions while the *zcf27Δ* strain grew similar to *Cg*2001 WT. Additionally, our studies demonstrated that Zcf27 and Zcf4 can associate with promoters of *ERG* genes, and their enrichment at these sites is further enhanced upon azole treatment or hypoxic conditions, respectively. Overall, we have discovered that *C. glabrata* maintains two Hap1 homologs to regulate ergosterol homeostasis. Specifically, Zcf27 aids in facilitating azole-mediated gene activation, while Zcf4 mediates hypoxia-induced gene repression.

## RESULTS

### Hap1 alters azole susceptibility in *S. cerevisiae*

In *S. cerevisiae*, there are three zinc cluster transcription factors Upc2, Ecm22 and Hap1 that are known to regulate the expression of ergosterol gene expression for sterol homeostasis (40–44). In addition, Upc2 and Ecm22 are also known to mediate azole susceptibility in *S. cerevisiae* (45, 46). However, until now, the role of Hap1 in altering azole susceptibility has not been determined. To test this hypothesis, the *hap1-Ty1* mutant was deleted in S288C strains BY4741 and FY2609 to generate BY4741 *hap1Δ* (this study) and FY2611 *hap1Δ* (40), respectively (see Supplemental Table S3). The indicated strains were tested for growth in liquid cultures and through serial-dilution spot assays with and without 16 μg/mL fluconazole (Fig. 1). Interestingly, the BY4741 strain exhibited a slight increase in fluconazole susceptibility compared to the BY4741 *hap1Δ* strain (Fig. 1A and C). We suspect that the enhanced azole susceptibility in the BY4741 strain is because of a known insertion of an in-frame *Ty1* sequence at the 3’ end of the *HAP1* ORF, resulting in the expression of a mutated *HAP1* that lacks 13 amino acids from its C-terminus and contains an additional 32 amino acids encoded from the *Ty1* sequence. The insertion of the *Ty1* element does not seem to affect the growth of BY4741 (*hap1-Ty1* mutant) versus FY2609 (*HAP1* WT) under untreated conditions (Fig. 1A-C, Table S1). In contrast, deletion of *HAP1* (FY2611 *hap1Δ*) showed a hypersusceptible phenotype compared to FY2609 when grown on agar plates or in liquid culture containing 32 μg/mL fluconazole (Fig. 1A and C). In addition, both the FY2611 *hap1Δ* strain and the BY4741 *hap1Δ* strain have a similar doubling time in the presence and absence of fluconazole (Fig. 1A-C and Table S1). To our knowledge, this is the first observation that Hap1 contributes to azole susceptibility in *S. cerevisiae*. We suspect that this phenotype has not been observed until now because earlier functional genomics screens used the BY4741 and BY4742 parental and deletion strain collections (47).

**FIG 1.**
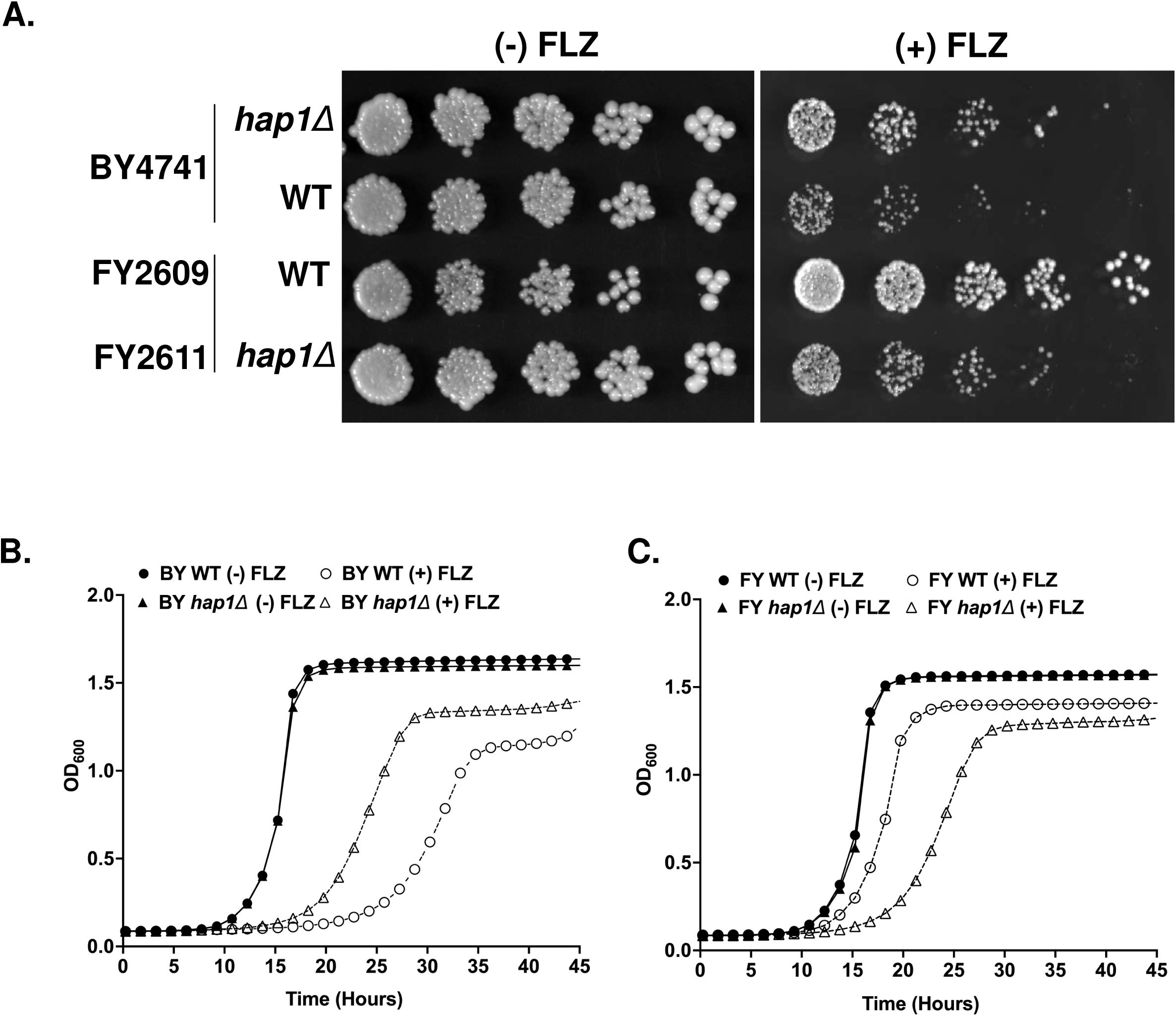
Zinc cluster transcription factor Hap1 in *S. cerevisiae* alters fluconazole susceptibility. **(A and B)** Fluconazole susceptibility of BY4741, BY4741 *hap1Δ*, FY2609, and FY2611 *hap1Δ* of *S. cerevisiae* S288C strains. Five-fold serial dilution assays of indicated strains grown on SC plates with and without 16 μg/ml fluconazole and incubated at 30°C for 48 hr. **(C and D)** Growth curve of indicated strains grown in SC liquid media with or without 16 μg/ml fluconazole.

### Phylogenetic analysis of Hap1 homologs in pathogenic fungi

A phylogenetic tree was constructed to investigate the evolutionary relationships of *S. cerevisiae* Hap1 homologs in the human pathogen *C. glabrata* and other fungal species. Two genes, Zinc cluster factor 4 (Zcf4) and Zinc cluster factor 27 (Zcf27), in *C. glabrata* grouped within the Hap1 clade of transcription factors (Fig. 2 and S1). *C. glabrata* is now recognized as a member of the *Nakaseomyces* genus (48). In our phylogeny, Zinc cluster factor 4 (Zcf4) and Zinc cluster factor 27 (Zcf27) in *C. glabrata* group with two non-pathogenic species of *Nakaseomyces*, *N. delphensis*, and *N. bacillisporus*. Although *Cg*Zcf4 groups more closely with *Sc*Hap1 in the tree compared to *Cg*Zcf27, support for the association is weak (ultrafast bootstrap support < 90). In general, the branching pattern of the Hap1 gene tree does not match the expected species relationships as determined by whole genome phylogenomic analysis (49). This suggests a complicated evolution history for this gene family including gene/genome duplication, gene loss, and possible horizontal gene transfer (50). The timing of the duplication event that gave rise to Zcf4 and Zcf27 in the *Nakaseomyces* clade is unclear. The duplication could have occurred in a common ancestor of *S. cerevisiae* and *C. glabrata,* and one copy was subsequently lost in *S. cerevisiae*. An alternative explanation is that the duplication occurred after the *Nakaseomyces-Saccharomyces* split.

**FIG 2.**
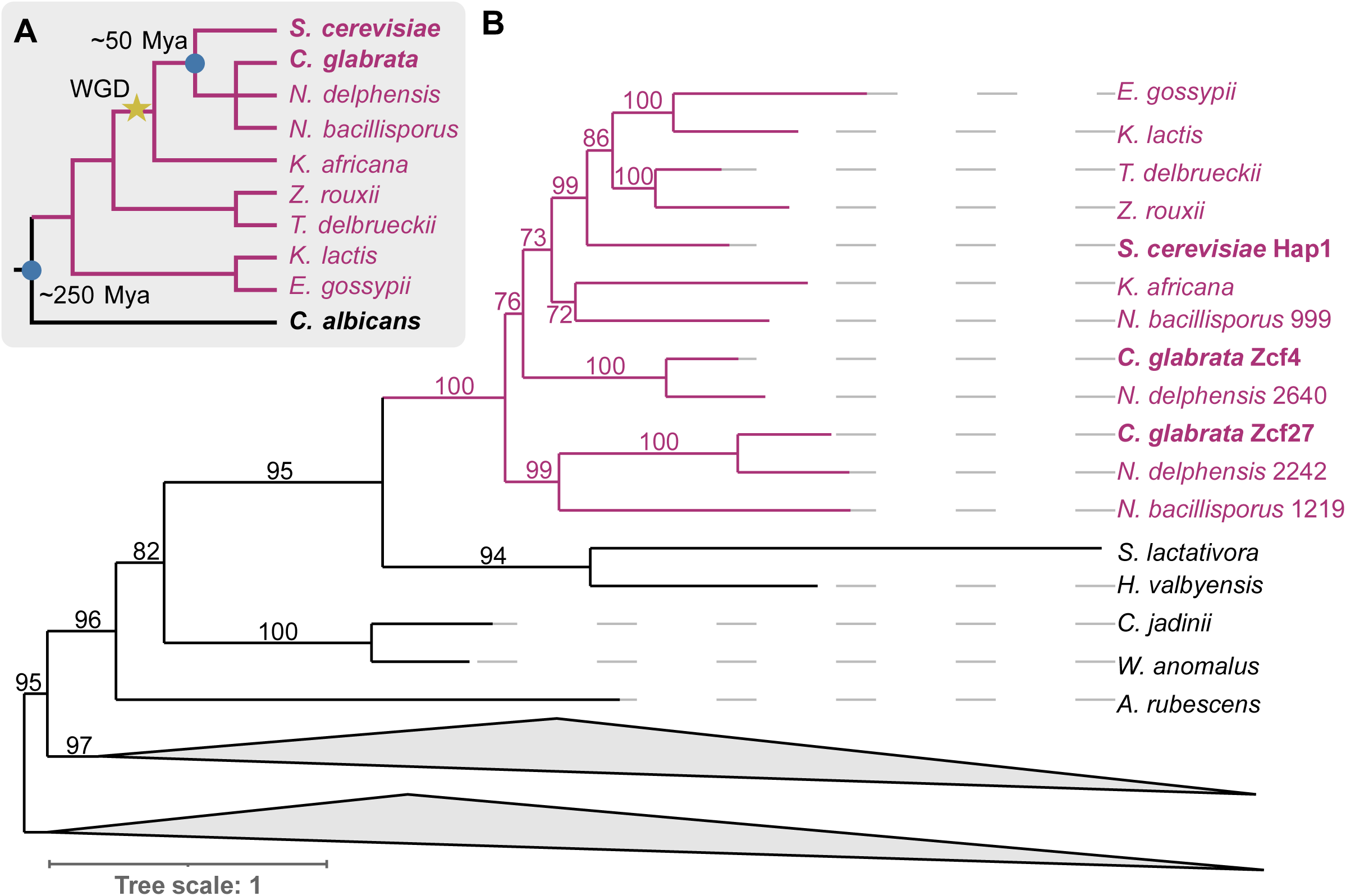
Evolutionary analysis of Hap1 homologs in fungi. **(A)** Species phylogeny showing the relationship of *C. albicans* and species within the *Saccharomycetaceae* (purple branches). Approximate divergence times of blue nodes are labeled. Location of the whole genome duplication event (WGD) is labeled. **(B)** Gene phylogeny of Hap1 homologs. The phylogeny was midpoint rooted and branch values represent ultrafast bootstrap support. Members of the *Saccharomycetaceae* are colored purple. Clades with more distantly related Hap1 homologs were collapsed for visualization (see Fig. S2 for the additional clades).

### Zcf27, rather than, Zcf4 alters azole susceptibility in *C. glabrata*

Because deletion of *HAP1* in *S. cerevisiae* altered azole susceptibility, we wanted to determine if *C. glabrata* strains lacking their Hap1 homologs Zcf27 and Zcf4 have a similar susceptibility to azole drugs. To test this hypothesis, we deleted *ZCF4* and *ZCF27* in the *C. glabrata* CBS138 (ATCC *Cg*2001) WT strain and performed liquid growth and serial-dilution spot assays with and without 32 µg/mL fluconazole (Fig. 3A-C). In the untreated conditions, both *zcf27Δ* and *zcf4Δ* strains grew similar to the *Cg*2001 WT strain on agar plates and liquid cultures (Fig. 3A-C). We also did not observe any differences in doubling times (Table S2). However, in the presence of fluconazole, the *zcf27Δ* strain showed an azole hypersusceptibility phenotype on agar plates along with a growth delay and longer doubling times when cultured in liquid media, whereas *zcf4Δ* strain grew like the *Cg*2001 WT strain (Fig. 3A and C, Table S2). To confirm that our observed azole hypersusceptible phenotype was due to the loss of *ZCF27*, the full-length *ZCF27* open-reading frame with its endogenous promoter were cloned in the pGRB2.0 plasmid and transformed into a *Cg*989 *zcf27Δ* deletion strain (Table S3 and S4). The pGRB2.0 vector was also transformed into *Cg*989 (ATCC 200989) as a control (Table S3 and S4). The *ZCF27* plasmid construct was able to rescue azole susceptibly as shown by a serial-dilution spot assay (Fig. 3D) while the *zcf27Δ* strain expressing the plasmid only construct remain hypersusceptible (Fig. 3D). In addition, gene expression analysis also confirmed that *ZCF27* and *ZCF4* were not expressed in their respective deletion strains (Fig. S2A and B). In addition, we confirmed that the genes upstream and downstream of *ZCF27* were expressed in *zcf27Δ* similar to the *Cg*2001 WT strain (Fig. S3A and B). Finally, we also deleted the upstream (*CAGL0K05819g*) and downstream (*CAGL0K05863g*) genes and observed no change in azole susceptibility (Fig. S3C). Overall, our data shows that Zcf27, rather than Zcf4, plays a specific role in mediating azole susceptibility.

**FIG 3.**
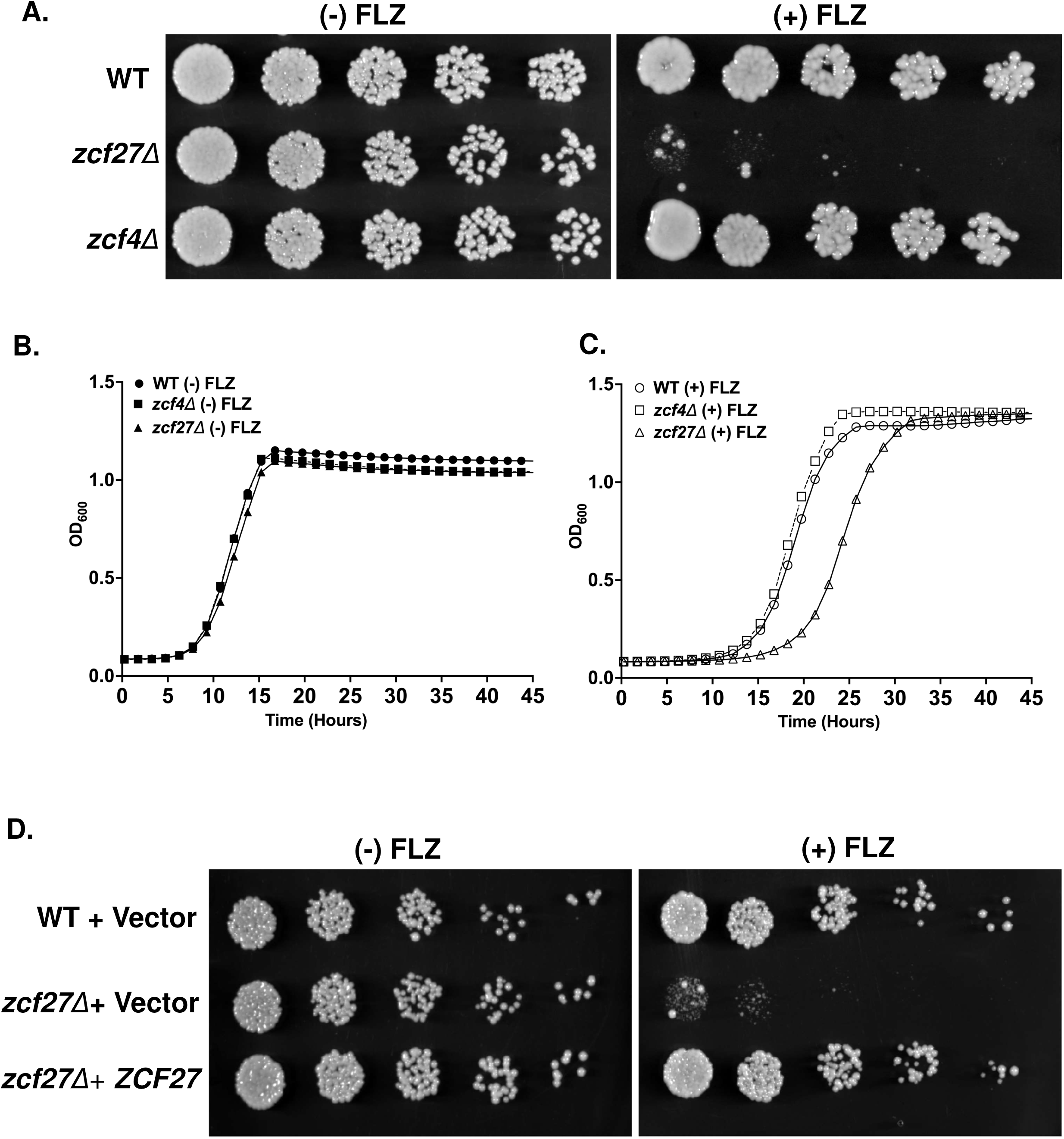
*C. glabrata* Zcf27, rather than, Zcf4 alters fluconazole susceptibility. **(A)** Five-fold serial dilution spot assays of *Cg*2001 WT, *zcf27Δ* and *zcf4Δ* strains plated on SC plates with and without 32 µg/mL fluconazole. **(B)** Liquid growth curves of the indicated *C. glabrata* strains grown in SC media with or without 32 µg/mL fluconazole. **(C)** Five-fold serial dilution assays of *Cg*989 WT and *zcf27Δ* transformed with plasmids expressing *ZCF27* from its endogenous promoter or empty vector spotted on SC-Ura plates with and without 32 µg/mL fluconazole at 30°C for 48 hr.

### Expression of *CYC1* depends on Zcf27, but not Zcf4 because of differences in protein expression

In *S. cerevisiae* Hap1 is known to regulate the expression of the *CYC1* gene (51–55). To determine if Zcf27 and/or Zcf4 also controls the expression of *C. glabrata CYC1* gene, *Cg*2001 WT, *zcf27Δ*, and *zcf4Δ* strains were grown in the presence and absence of azole treatment and qRT-PCR transcript analysis was performed. Interestingly, *CYC1* transcript analysis revealed that the loss of *ZCF27*, but not *ZCF4*, resulted in a 50% decrease in *CYC1* expression, irrespective of drug treatment (Fig. 4A and B). To determine if this difference was a consequence of transcript levels of *ZCF4* and *ZCF27,* qRT-PCR analysis was performed on *Cg*2001 WT cells treated with or without 64 µg/mL fluconazole for 3 or 6 hours. Both *ZCF27* and *ZCF4* transcript levels were expressed with no significant differences between untreated and fluconazole treated conditions (Fig. 4C and D; Table S7). Furthermore, *ZCF27* transcript levels are not altered in *zcf4Δ* strain and vice versa indicating they are independent of each other (Fig. S2A and B). To determine if protein expression levels differed between Zcf27 and Zcf4, we constructed endogenously 3XFLAG tagged strains where the 3XFLAG tag was inserted at the C-terminus of *ZCF27* and *ZCF4*. After PCR confirmation, Zcf27-3XFLAG and Zcf4-3XFLAG tagged strains were grown with or without 64 µg/mL fluconazole for 3 or 6 hrs. Western blot analysis indicated that the Zcf27-3XFLAG protein expression remained fairly constant with and without drug treatment (Fig. 4E). Unexpectedly, we observed virtually no expression of Zcf4-3XFLAG protein regardless of drug treatment (Fig. 4E, Short Exp). Even with longer exposure times, barely detectable levels of Zcf4 were observed (Fig. 4E, Long Exp) suggesting that Zcf4 is regulated at the post-transcriptional level. Due to essentially undetectable levels of Zcf4 protein, we suspect that this is why a *zcf4Δ* strain does not alter *CYC1* gene expression or show hypersusceptibility to azoles.

**FIG 4.**
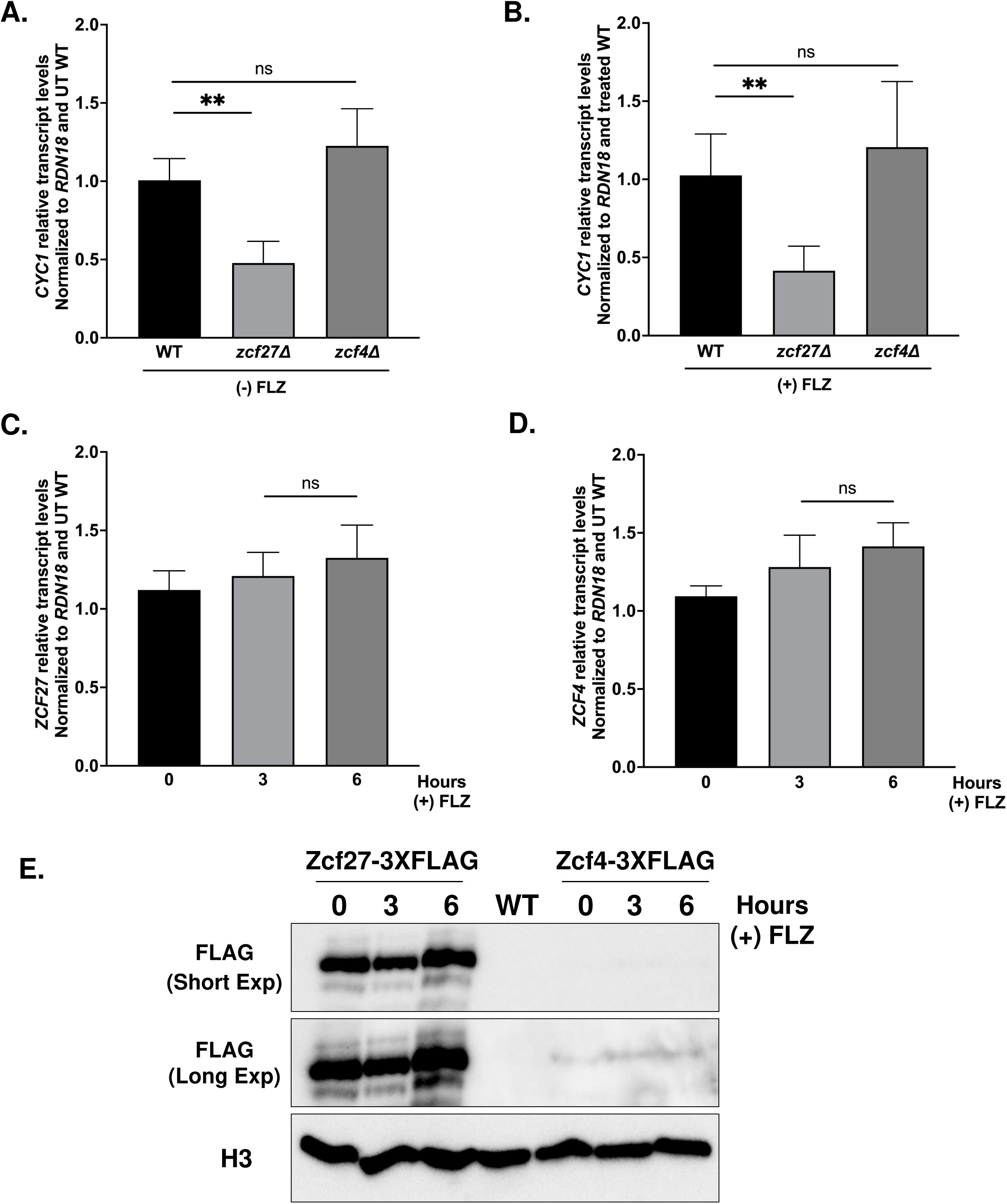
Transcript and protein analysis of *CYC1*, *ZCF27* and *ZCF4*. **(A and B)** Transcript levels of *CYC1* from the indicated strains treated with and without 64 µg/mL fluconazole for 3 hr. **(C and D)** Transcript levels of *ZCF27* and *ZCF4* from the indicated strains treated with and without 64 µg/mL fluconazole for 3 hr. For A-D, transcript levels were set relative to WT and normalized to *RDN18* mRNA levels. Data were analyzed from three or more biological replicates with three technical replicates. Statistics were determined using the GraphPad Prism Student t-test, version 9.5.1. Error bars represent SD. ns, P > 0.05; **P < 0.01. **(E)** Western blot analysis of Zcf27-3X FLAG and Zcf4-3XFLAG with and without treatment with 64 µg/mL fluconazole for 3 hr and 6 hr. Western blots showing short (Short Exp) and long (Long Exp) enhanced chemiluminescence exposure. Histone H3 was used as the loading control.

### Zcf27 is dispensable for expression of drug efflux pumps but is needed for azole-induced expression of ergosterol (*ERG*) genes

Because the *zcf27Δ* strain showed altered azole susceptibility (Fig.3A-C), we wanted to identify the mechanism mediating this phenotype. A common mechanism of altering azole resistance in *C. glabrata* involves the upregulation of drug efflux pumps such as *CDR1*, *PDH1*, and *SNQ2*, facilitated by the zinc cluster transcription factor Pdr1 (28, 29, 31, 56, 57). To determine if expression of drug efflux pumps is altered in the *zcf27Δ* strain in the presence or absence of 64 µg/mL fluconazole, the expression levels of the known azole transporters *CDR1, PDH1,* and *SNQ2* as well as the transcriptional regulator *PDR1* were analyzed by qRT-PCR analysis. Our transcript analysis revealed no significant difference in the expression of any of the genes encoding ABC-transporters in the *zcf27Δ* strain compared to the *Cg*2001 WT strain (Fig. 5A and B; S4A and B) indicating that altered expression of azole drug efflux pumps is not the reason for azole hypersusceptibility for the *zcf27Δ* strain.

**FIG 5.**
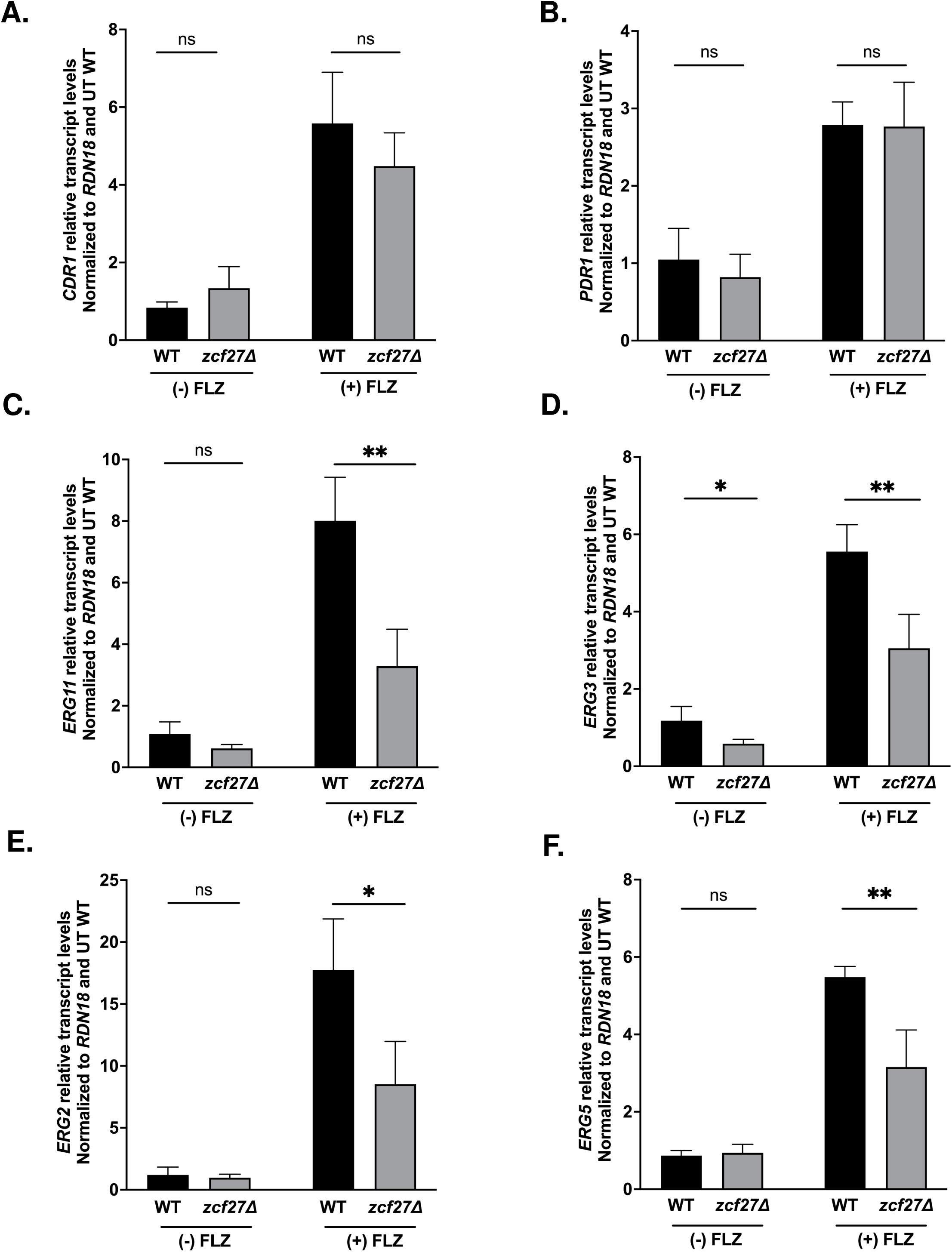
Transcript analysis of drug efflux pump and ergosterol (*ERG*) genes. **(A and B)** Transcript levels of drug efflux pumps in *Cg*2001 WT and *zcf27Δ* strains treated with or without 64 µg/mL fluconazole for 3 hr. **(C-F)** Transcript levels of *ERG* genes in *Cg*2001 WT and *zcf27Δ* strains treated with or without 64 µg/mL fluconazole for 3 hr. For A-F, all strains were treated with or without 64 µg/mL fluconazole for 3 hr. Transcript levels were set relative to the untreated WT and normalized to *RDN18* mRNA levels. Data were analyzed from four biological replicates with three technical replicates each. Statistics were determined using the GraphPad Prism Student t test, version 9.5.1. ns, P > 0.05; *, P < 0.05; **, P <0.01. Error bars represent the SD.

In *S. cerevisiae*, Hap1 is known to regulate steady state transcript levels of ergosterol biosynthesis genes such as *ERG11*, *ERG3*, *ERG5* and *ERG2* (40, 41, 44, 52, 53, 58, 59) . In addition, altered *ERG11* gene expression in *C. glabrata* is also a mechanism that can lead to azole hypersusceptibility phenotypes (24, 60, 61). To determine if altered *ERG* gene expression was a mechanism for the observed azole hypersusceptibility of the *zcf27Δ* strain, *Cg*2001 WT and *zcf27Δ* strains were treated with and without 64 µg/mL fluconazole and *ERG11*, *ERG3*, *ERG5* and *ERG2* transcript levels were analyzed by qRT-PCR. In the absence of drug, with the exception of *ERG3*, no significant difference in the expression levels of *ERG11*, *ERG5* or *ERG2* was observed between the *Cg*2001 WT and *zcf27Δ* strain (Fig. 5C-F). However, upon treatment with fluconazole, all four *ERG* genes failed to induce to wild-type levels in the *zcf27Δ* strain (Fig. 5C-F). Furthermore, a *zcf4Δ* strain did not have altered *ERG11* and *ERG3* expression which coincides with its lack of expression and azole hypersusceptible phenotype (Fig. S4C and D). Altogether, our data indicates that in addition to Upc2A, Zcf27 serves as another critical transcription factor for the azole-induced expression of the late ergosterol pathway genes.

### Zcf27-3XFLAG is enriched at *ERG* gene promoters

Because our data shows decreased expression of ergosterol genes in the *zcf27Δ* strain upon azole treatment (Fig. 5C-F), we suspect that Zcf27 is a direct transcription factor for the *ERG* genes. To determine if Zcf27 directly targets the promoter of the *ERG11* gene, chromatin immunoprecipitation (ChIP) assays were performed using anti-FLAG monoclonal antibodies and chromatin isolated from untagged *Cg*2001 WT and Zcf27-3XFLAG strains, treated with or without fluconazole. ChIP-qPCR fluorescent probes were designed to recognize a distal (E11P1) and proximal (E11P2) promoter region of *ERG11*. Using these probes, a significant enrichment of Zcf27 was detected at both *ERG11* promoter regions compared to the untagged control (Fig. 6A and B; Table S8). In addition, Zcf27 was further enriched at the promoter of *ERG11* upon azole treatment (Fig. 6A and B; Table S8) supporting the importance of Zcf27 in azole-induced gene expression. No significant enrichment of Zcf27 was detected at the 3’*UTR* of *ERG11* regardless of treatment (Fig. S5), indicating specific enrichment at the promoter region.

**Fig 6.**
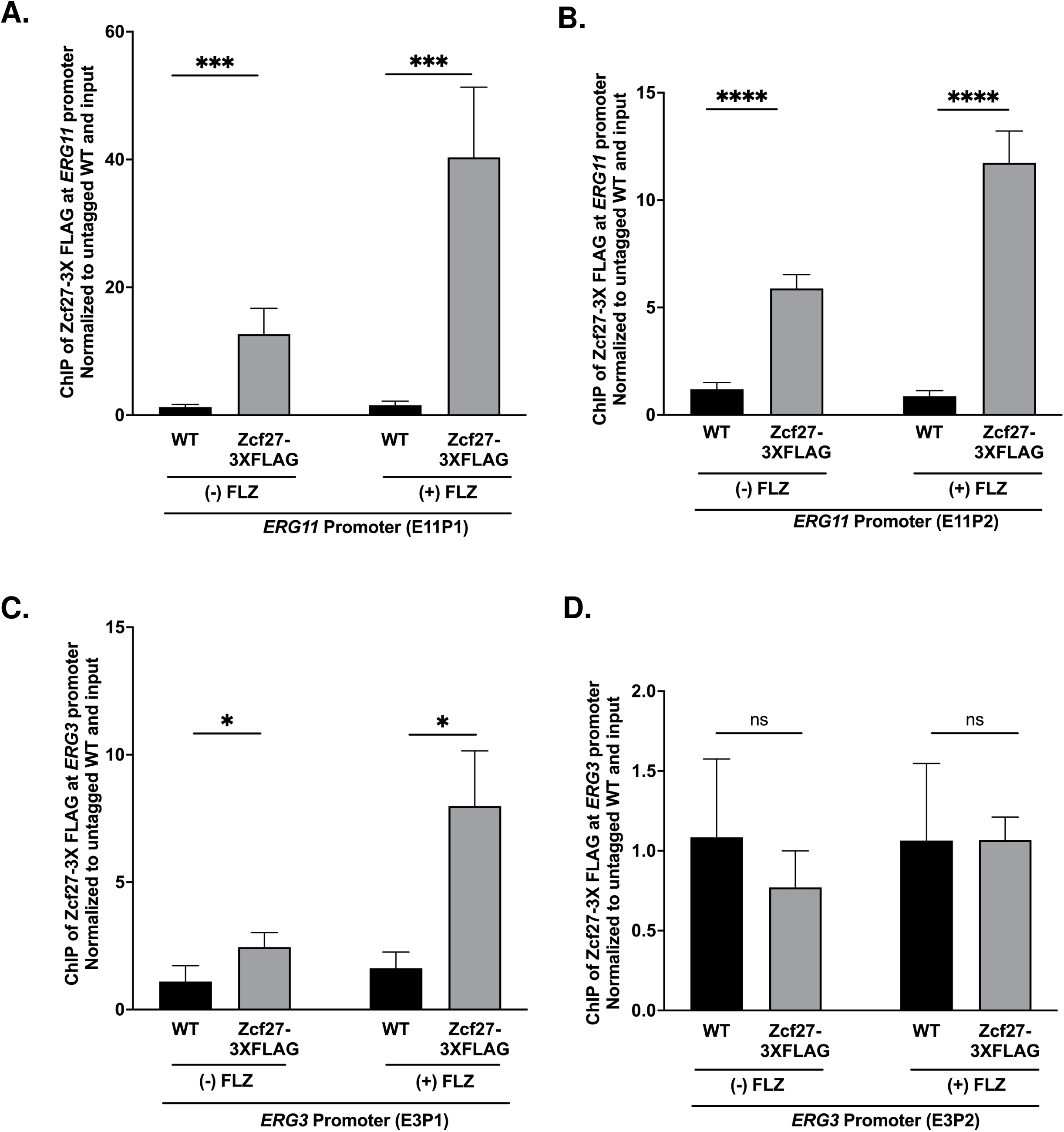
Chromatin immunoprecipitation analysis of Zcf27 at *ERG* gene promoters. **(A and B)** ChIP analysis of *Cg*2001 WT (untagged) and Zcf27-3XFLAG at two *ERG11* promoter regions (E11P1 and E11P2) when treated with or without 64 µg/mL fluconazole for 3 hr. **(C and D)** ChIP analysis of Zcf27-3XFLAG at two *ERG3* promoter regions (E3P1 and E3P2) when treated with and without 64 μg/mL fluconazole for 3 hr. For A-D, ChIP analysis was normalized to DNA input samples and set relative to untagged WT. Statistics were determined using the GraphPad Prism Student t test, version 9.5.1. ns, P > 0.05; *, P < 0.05; ***, P < 0.001; ****, P < 0.0001. Error bars represent SD for three biological replicates with three technical replicates.

We also examined Zcf27 localization status on the *ERG3* promoter by ChIP analysis (Fig 6C and D). To determine if Zcf27 binds to the promoter of *ERG3,* two ChIP-qPCR fluorescent probes were designed to recognize the distal (E3P1) and proximal (E3P2) promoter regions.

Similar to the *ERG11* promoter, Zcf27 was detected at the distal *ERG3* promoter region and was further enriched upon fluconazole treatment (Fig. 6C and Table S8). However, we did not detect any Zcf27 enrichment at the more proximal promoter region (Fig. 6D and Table S8) regardless of azole treatment. Overall, our data demonstrates that Zcf27 directly targets the promoters of *ERG11* and *ERG3* to help facilitate the proper expression of *ERG* genes and maintenance of ergosterol homeostasis during azole treatment.

### The *zcf27Δ* strain has altered azole susceptibility due to decreased ergosterol levels, which can be suppressed by exogenous sterols and active sterol import

Because azole-induced *ERG* gene expression is diminished in the *zcf27Δ* strain, we would expect an additional decrease in ergosterol levels in this strain, which would explain why a *zcf27Δ* strain has an increase in azole susceptibility. To ascertain whether total endogenous ergosterol levels differed between *Cg*2001 WT and *zcf27Δ* strains upon azole treatment, non-polar lipids were extracted from both strains in the presence and absence of 64 µg/mL fluconazole. Total ergosterol level was measured by high performance liquid chromatography (HPLC) analysis and cholesterol was used as an internal standard control. No significant difference was observed between *Cg*2001 WT and *zcf27Δ* strains in the untreated conditions, concurring with our gene expression analysis showing no significant difference in expression of multiple *ERG* genes without azole treatment (Fig. 7A). However, upon fluconazole treatment, the *Cg*2001 WT strain demonstrated the expected decrease in ergosterol levels (Fig. 7B), whereas the *zcf27Δ* strain exhibited an additional 30% reduction in total ergosterol compared to the treated *Cg*2001 WT strain (Fig. 7C).

**Fig 7.**
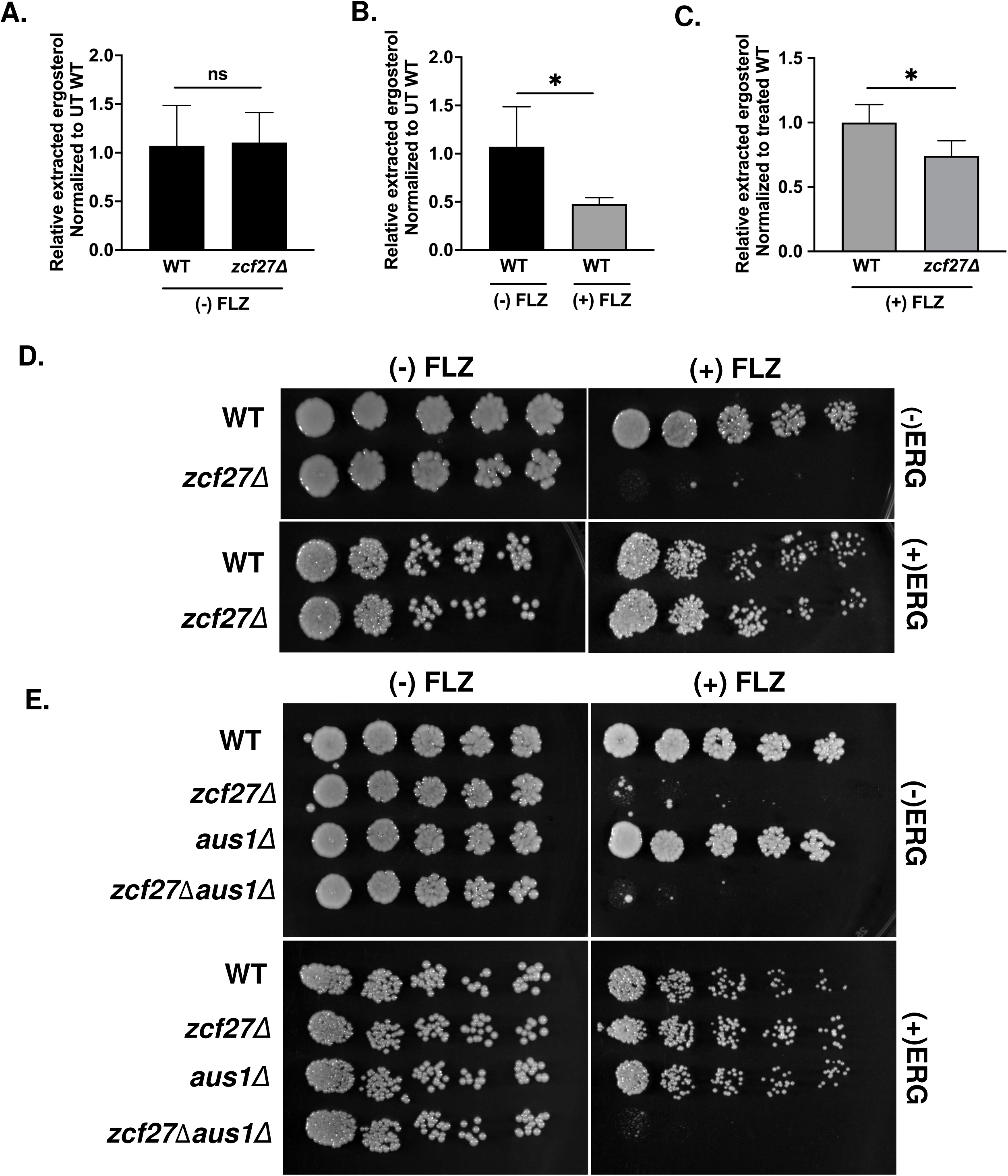
Disruption of ergosterol levels in a *zcf27Δ* strain alters azole susceptibility **(A-C)** HPLC analysis of the total ergosterol extracted from the *Cg*2001 WT and *zcf27Δ* strains treated with or without 64 µg/mL fluconazole for 3 hr. The figure represents a ratio between ergosterol and cholesterol, compared to treated and untreated WT samples. Data were generated from four biological replicates. Statistics were determined using the GraphPad Prism Student t test, version 9.5.1. ns, P > 0.05; *, P < 0.05. Error bars represent the SD. **(D and E)** Five-fold dilution spot assays of the indicated *C. glabrata* strains grown on SC plates with and without 32 µg/mL fluconazole and/or with and without 20 µg/mL ergosterol.

Due to this observation, we hypothesized that the decrease in ergosterol content contributes to azole hypersensitivity and reasoned that exogenous supplementation with ergosterol would suppress the azole hypersensitive phenotype observed for the *zcf27Δ* strain. To test this hypothesis, *Cg*2001 WT and *zcf27Δ* strains were plated on synthetic complete (SC) media supplemented with or without exogenous ergosterol and/or fluconazole. In support of our hypothesis, serial-dilution spot assays showed that the addition of exogenous ergosterol completely suppressed the azole hypersensitive phenotype of the *zcf27Δ* strain, whereas *zcf27Δ* strain without ergosterol retained the hypersensitive phenotype (Fig. 7D). Because ergosterol is solubilized in the presence of Tween 80-ethanol solution, we wanted to determine if this suppression was specific to ergosterol. Thus, *Cg*2001 WT and *zcf27Δ* strains were plated on SC media supplemented with a Tween 80-ethanol solution with or without fluconazole. As indicated in supplemental Fig. S6, Tween 80-ethanol did not suppress *zcf27Δ* azole hypersusceptible phenotype (Fig. S6) indicating that suppression was mediated by exogenous ergosterol uptake.

Based on these observations, we also expected that deletion of the only known sterol importer *AUS1* would prevent sterol uptake by *zcf27Δ* strains (62–64). To determine this, we constructed an *aus1Δ* strain and a *zcf27Δaus1Δ* double deletion strain and performed serial-dilution spot assays on agar plates supplemented with or without exogenous ergosterol in the presence and/or absence of fluconazole (Fig. 7E). As anticipated, the *zcf27Δaus1Δ* strain remained hypersensitive to fluconazole with or without exogenous ergosterol (Fig. 7E).

However, growth of the *aus1Δ* strain was not altered by fluconazole and/or exogenous ergosterol and grew similar to the *Cg*2001 WT strain (Fig. 7E). Overall, our data elucidates the mechanistic basis and pathway underlying the hypersensitive phenotype observed in the *zcf27Δ* strain. Because Zcf4 is not expressed, it is unclear what role it plays, if any, under azole treatment. In summary, our findings represent the first characterization of Zcf27 as direct transcription factor for regulating ergosterol genes and ergosterol homeostasis in response to azole drug treatment.

### Zcf4 is induced upon hypoxic exposure

In aerobic conditions, *S. cerevisiae* Hap1 functions as a transcriptional activator of *CYC1* and *ERG* genes (40, 41, 44, 51–55, 58, 59). Furthermore, our presented data suggests that Zcf27 operates similarly to Hap1, by regulating the corresponding conserved genes in *C. glabrata*. Interestingly, in *S. cerevisiae*, Hap1 functions also as a transcriptional repressor to shut down *ERG* genes under hypoxia by recruiting a corepressor complex containing Set4, Tup1, and Ssn6 corepressors (40, 59, 65, 66). Currently, it is not known if Zcf27, Zcf4 or another transcription factor functions to repress *C. glabrata ERG* genes under hypoxic conditions.

Due to our observed phenotype for the *zcf27Δ* strain, but not for the *zcf4Δ* strain under azole treated conditions, *C. glabrata Cg*2001 WT, *zcf27Δ*, and *zcf4Δ* strains were serially diluted five-fold on agar plates and grown under aerobic or hypoxic conditions (Fig. 8A). Interestingly, under hypoxic conditions, only the *zcf4Δ* strain exhibited a statistically significant slow growth defect, as determined by colony size (Fig. 8A and B). Measuring the colony diameter revealed an approximate 40% decrease in the size of *zcf4Δ* colonies when compared to both the *Cg*2001 WT and the *zcf27Δ* colonies suggesting a potential function for Zcf4 (Fig. 8B). Due to the significant differences in protein expression observed between Zcf27 and Zcf4 under aerobic conditions, we also evaluated the transcript and protein expression levels of Zcf4 and Zcf27 under hypoxic conditions. Using qRT-PCR analysis, a 4-fold increase in *ZCF4* transcript levels was detected after two hours under hypoxic conditions while *ZCF27* transcript levels remained unaltered from aerobic to hypoxic conditions (Fig. 8C and D). In addition, we assessed the protein levels of Zcf4-3XFLAG and Zcf27-3XFLAG tagged strains using Western blot analysis. Remarkably, we detected robust levels of Zcf4 proteins under hypoxic conditions while Zcf27 protein levels remained the same from aerobic to hypoxic conditions (Fig. 8E and F). Taken together, we have identified Zcf4 as the first hypoxia-inducible transcription factor in *C. glabrata*. Given that *S. cerevisiae* Hap1 is required for repressing *ERG* genes under hypoxic conditions, we anticipate that Zcf4 is hypoxia-induced to function in a similar manner.

**Fig 8.**
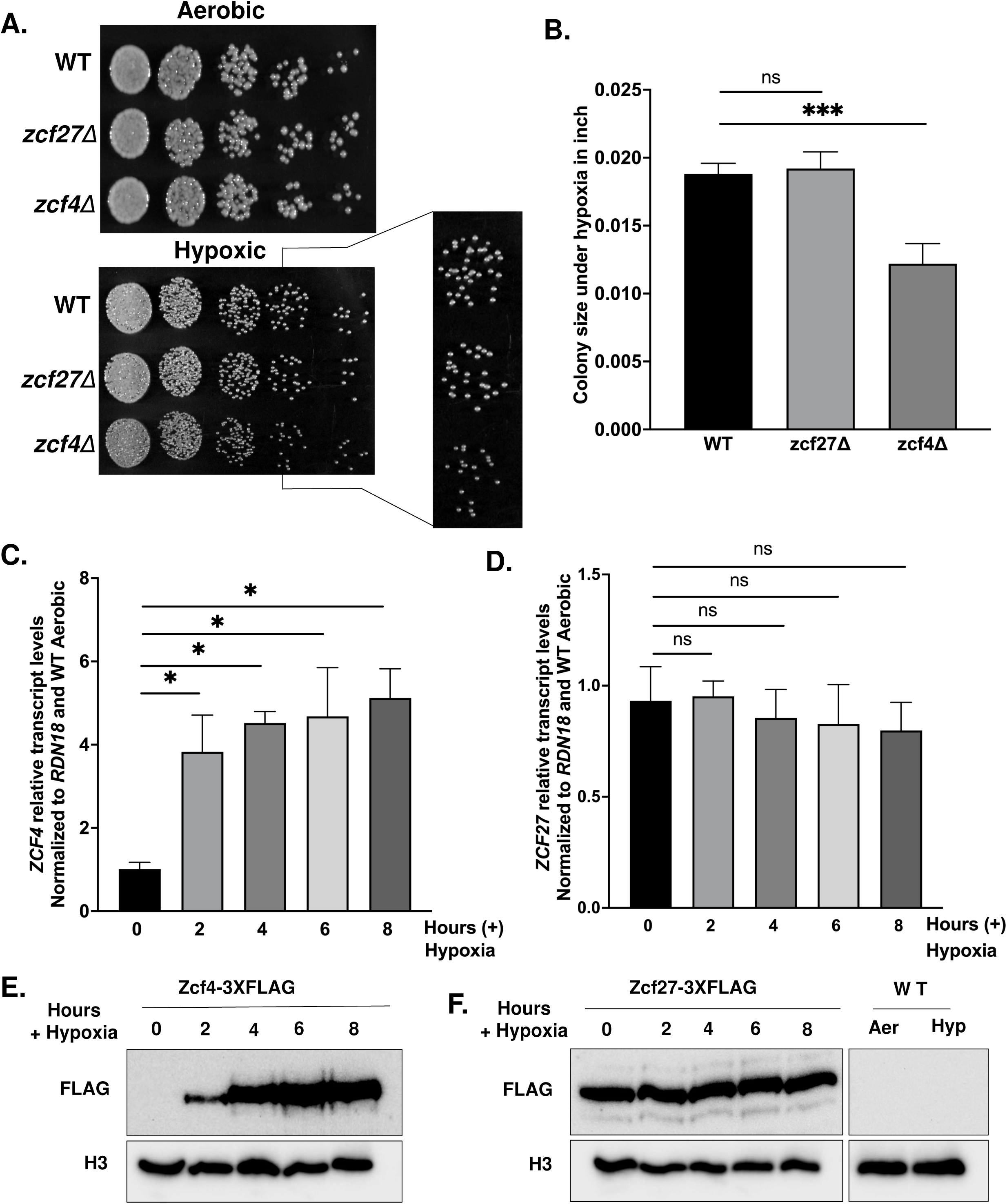
Phenotypic and expression analysis of *C. glabrata* strains under hypoxic conditions **(A)** Five-fold serial dilution spot assays of *Cg*2001 WT, *zcf27Δ* and *zcf4Δ* strains grown on YPD plates under aerobic and hypoxic condition. The fourth dilution of the hypoxic plate was enlarged for enhanced visibility **(B)** Graphical representation of colony sizes of the indicated strains when grown under hypoxic conditions. Colony sizes were measured using ImageJ, version 1.51. Statistics were determined using the GraphPad Prism Student t test, version 9.5.1. ns, P > 0.05; ***, P < 0.001. **(C and D)** Transcript analysis of *ZCF4* and *ZCF*27 of the *Cg*2001 WT strain when grown under hypoxic conditions over a time course of 0, 2, 4, 6, and 8 hrs. The relative transcript levels were set to *Cg*2001 WT before hypoxic exposure (0 hr) and normalized to *RDN18*. Statistics were determined using the GraphPad Prism Student t test, version 9.5.1. ns, P > 0.05; *, P < 0.05. **(E and F)** Western blot analysis of Zcf4-3XFLAG and Zcf27-3XFLAG protein levels over a time course of 0, 2, 4, 6, and 8 hr of hypoxic exposure. *Cg*2001 WT (untagged) strain was used as a negative control for both aerobic (Aer) and hypoxic (Hyp) conditions. Histone H3 was used as a loading control.

### Ergosterol genes are downregulated upon hypoxic conditions

In *S. cerevisiae,* it is well established that exposure to hypoxia leads to the repression of the *ERG* pathway (40, 59, 65). To determine if hypoxia-mediated repression of *ERG* genes is conserved and robust in *C. glabrata*, as observed in *S. cerevisiae*, we performed transcript analysis of multiple *ERG* genes involved in the late ergosterol biosynthesis pathway, namely, *ERG11*, *ERG3, ERG2*, *ERG5*. When comparing the indicated *ERG* gene transcript levels under aerobic versus hypoxic conditions, we observed a significant decrease of 70-90% in expression under hypoxic conditions (Fig. 9A-D). These findings confirm that a conserved mechanism between *S. cerevisiae* and *C. glabrata* is maintained for shutting down ergosterol biosynthesis in response to hypoxic conditions.

**Fig 9.**
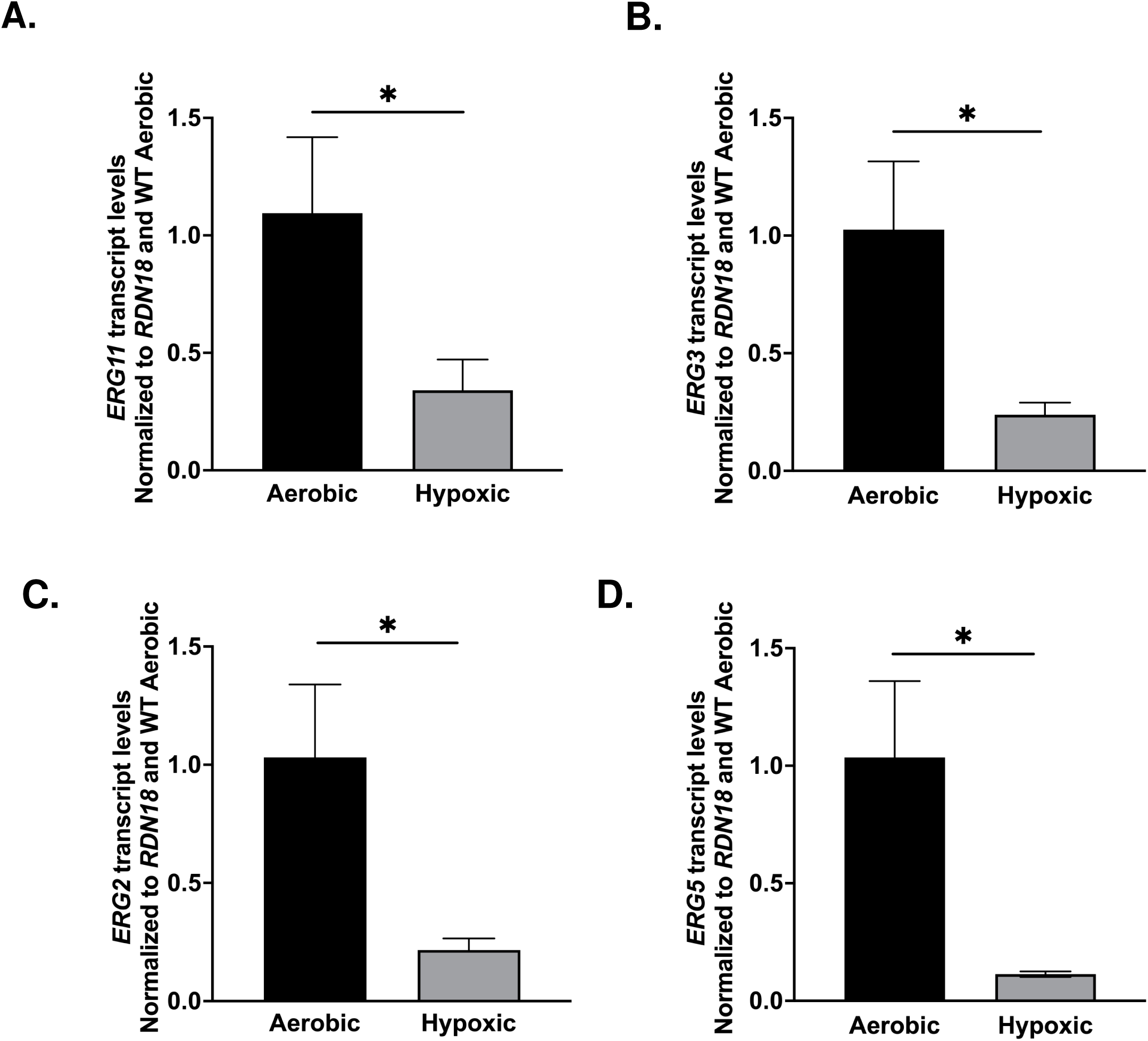
Ergosterol biosynthesis genes in *C. glabrata* are repressed upon hypoxic exposure. **(A-D)** The expression of *ERG11*, *ERG3*, *ERG2*, and *ERG5* was analyzed in *C. glabrata* WT cells under both aerobic and hypoxic conditions. Transcript analysis was set relative to aerobic WT and normalized to *RDN18* as the internal control. Data were collected from a minimum of three biological replicates, each with three technical replicates. Statistics were determined using the GraphPad Prism Student t test, version 9.5.1. ns, P > 0.05; *, P < 0.05. Error bars represent the SD.

### Zcf4, rather than Zcf27, represses genes from ergosterol pathway under hypoxic conditions

In *S. cerevisiae,* it is known that following exposure to hypoxia *ERG* genes are repressed by a WT copy of *HAP1* but not by *hap1-Ty1* expressed in S288C strains (40, 59, 65). To determine if Zcf27 and/or Zcf4 shares the same function as Hap1 under hypoxic conditions, qRT-PCR analysis on *ERG* genes were performed. Surprisingly, our transcript analysis did not detect any significant differences in the transcript levels of *ERG11*, *ERG3*, *ERG5* and *ERG2* between the *Cg*2001 WT and *zcf27Δ* strain under hypoxic conditions (Fig. 10A-D). In contrast, we observed a significant increase in the transcript levels of *ERG11*, *ERG3*, and *ERG5* genes in the *zcf4Δ* compared to *Cg*2001 WT strain (Fig. 10 E-G). Interestingly, *ERG2* showed no significant difference in the transcript levels upon hypoxic exposure in either *zcf27Δ* or *zcf4Δ* strain (Fig. 10D and H), despite being repressed upon hypoxic exposure (Fig. 9C) indicating involvement of another transcription factor. Overall, our findings suggest that Zcf4, rather than Zcf27, is directly or indirectly involved in hypoxia-induced *ERG* gene repression.

**Fig 10.**
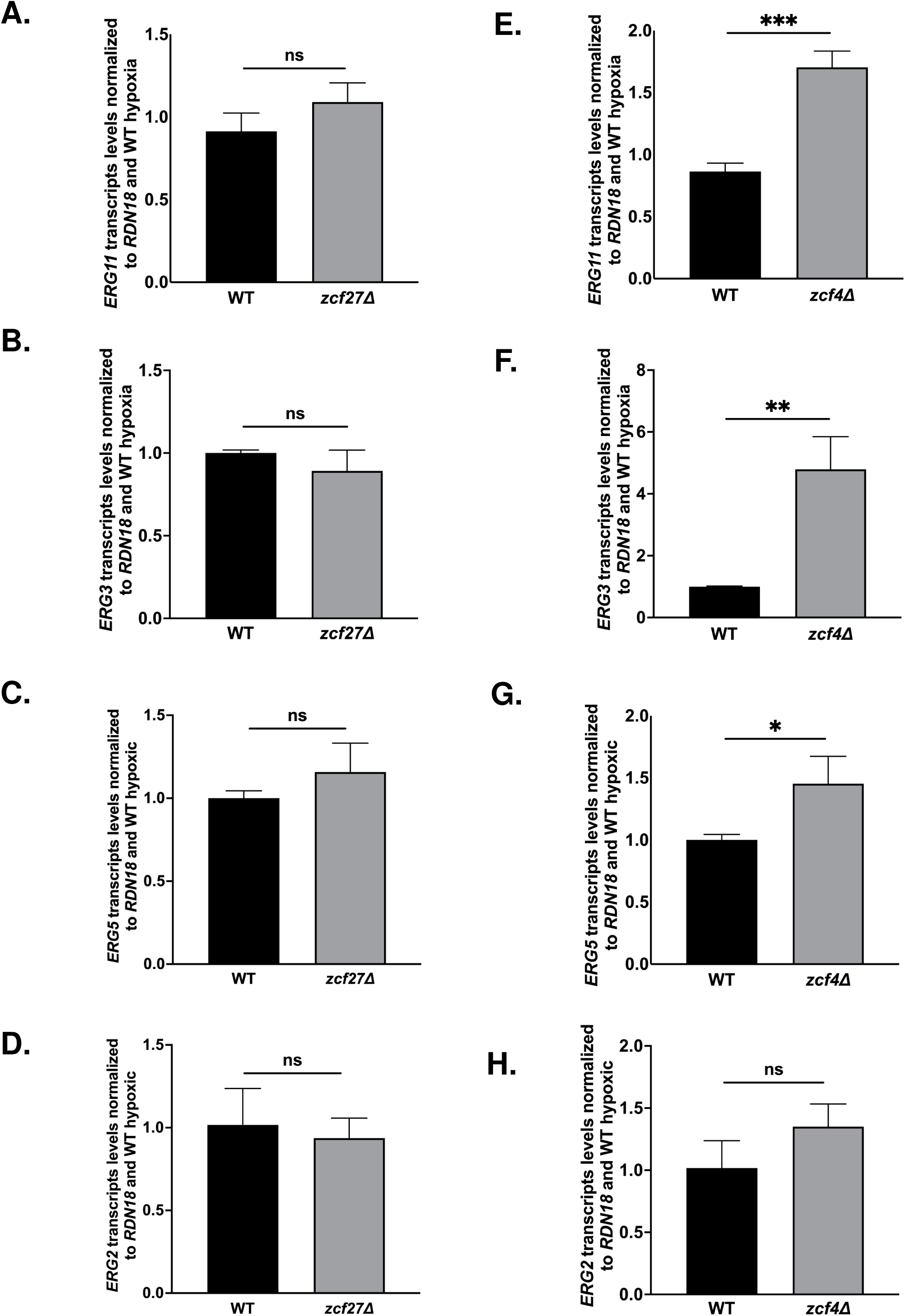
*ERG* genes are repressed by Zcf4 rather than Zcf27 upon hypoxic conditions. **(A-E)** After 8 hr of hypoxic exposure, transcript levels of *ERG11*, *ERG3*, *ERG5* and *ERG2* from the indicated strains were determined by qRT-PCR analysis. The levels of *ERG* genes were set relative to WT and normalized to *RDN18*. Data were generated from a minimum of three biological replicates. Statistics were determined using the GraphPad Prism Student t test, version 9.5.1. ns, P > 0.05; *, P < 0.05; **, P <0.01; ***, P < 0.001. Error bars represent the SD.

### Both Zcf4-3XFLAG and Zcf27-3XFLAG are enriched on *ERG11* and *ERG3* gene promoter upon hypoxic exposure

Because we determined that Zcf27 was enriched at the promoter sequences of *ERG11* and *ERG3* under aerobic azole conditions, we wanted to assess the direct binding of Zcf27 and Zcf4 at *ERG* gene promoters under hypoxic conditions. To determine this, ChIP assays were performed using anti-FLAG monoclonal antibodies and chromatin isolated from untagged *Cg*2001 WT, Zcf27-3XFLAG and Zcf4-3XFLAG strains grown for 8 hours under hypoxic conditions. The same ChIP-qPCR fluorescent probes used under azole-treated conditions were utilized to assess the enrichment of Zcf27 and Zcf4 at the *ERG11* and *ERG3* promoters. At the *ERG11* promoter, Zcf27 showed 3.5-fold enrichment at the proximal promoter sequence but was not enriched at the more distal promoter sequence (Fig. 11A and B). Interestingly, this differs from our observations under azole treated conditions, where Zcf27 was more enriched at the distal promoter sequence than the more proximal promoter sequence (Fig. 6A and B). For Zcf4, we observed a 5-fold enrichment at the *ERG11* distal promoter sequence compared to untagged *Cg*2001 WT strain, but no enrichment was observed at the proximal promoter sequence (Fig. 11C and D). In addition, Zcf27 and Zcf4 enrichment was specific to the promoter of *ERG11* since no significant enrichment was observed at the 3’*UTR* of *ERG11* (Fig. S7A and B). At the *ERG3* promoter, Zcf27 showed a 3-fold enrichment at the proximal promoter sequence but was not enriched at the distal promoter sequence (Fig. 11E and F). Again, this differs from our observations under azole treated conditions where Zcf27 enriches exclusively at the *ERG3* distal promoter sequence but not at the proximal promoter sequence (Fig 6C and D). In contrast, under hypoxic conditions, Zcf4 was 2-fold enriched at the *ERG3* distal promoter sequence but 20-fold enriched at the proximal promoter sequence suggesting the Zcf4 occupies both sites but prefers the more proximal sequence (Fig. 11G and H). Based on our observations, Zcf4 binding at these promoters likely prevents efficient binding of Zcf27 and Upc2A under hypoxic conditions so that *ERG* gene repression can occur.

**Fig 11.**
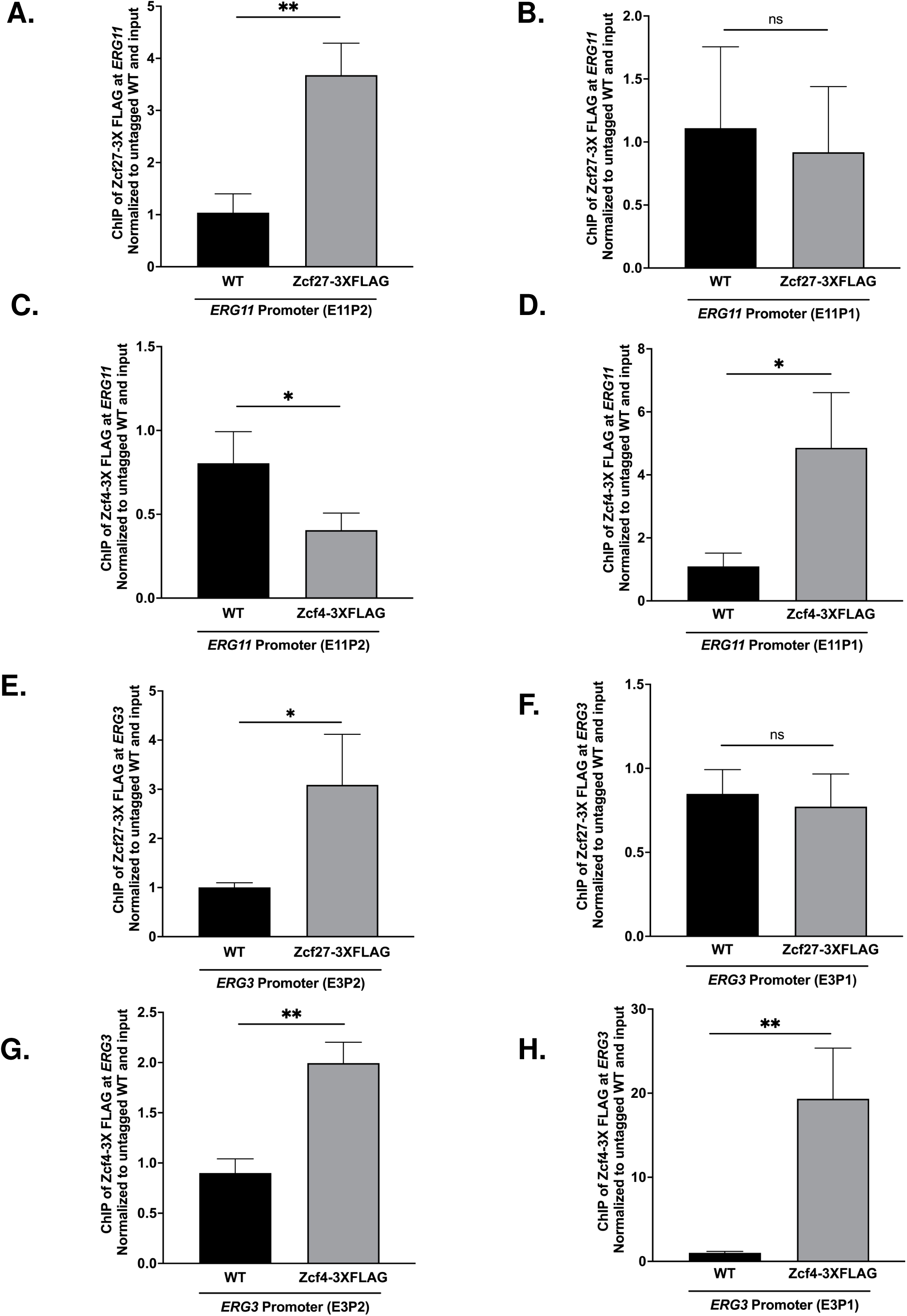
Chromatin immunoprecipitation analysis of Zcf27 and Zcf4 at *ERG* gene promoters under hypoxic conditions. **(A-H)** ChIP analysis of *Cg*2001 WT (untagged), Zcf27-3XFLAG and Zcf4-3XFLAG at two *ERG11* promoter regions (E11P1 and E11P2) and two *ERG3* promoter regions (E3P1 and E3P2) after 8 hr of hypoxic treatment. For A-H, ChIP analysis was normalized to DNA input samples and set relative to untagged WT. Data were generated from three biological replicates, with three technical replicates each. Statistics were determined using the GraphPad Prism Student t test, version 9.5.1. ns, P > 0.05; *, P < 0.05; **, P <0.01. Error bars represent the SD.

## DISCUSSION

In this study, the roles of the *S. cerevisiae* Hap1 zinc cluster transcription factor homologs, Zcf27 and Zcf4, were investigated in response to azole drug treatment and hypoxic conditions. Our data suggest that Zcf27 functions similarly to *Sc*Hap1 under aerobic conditions, regulating the conserved genes *CYC1* and *ERG3* under untreated conditions. Additionally, we found that loss of *ZCF27*, but not *ZCF4*, impacts azole susceptibility due to the inability to adequately induce *ERG* genes under azole drug treatment and maintain ergosterol homeostasis. Furthermore, we discovered that Zcf4 is specifically expressed in response to hypoxia, allowing it to function as a repressor of *ERG* genes. Overall, our study revealed that *C. glabrata* maintains two Hap1 homologs, Zcf27 and Zcf4, to control gene expression and mediate proper ergosterol homeostasis in response to both azole drug treatment and hypoxic conditions (see model Fig. 12A and B).

**Fig 12.**
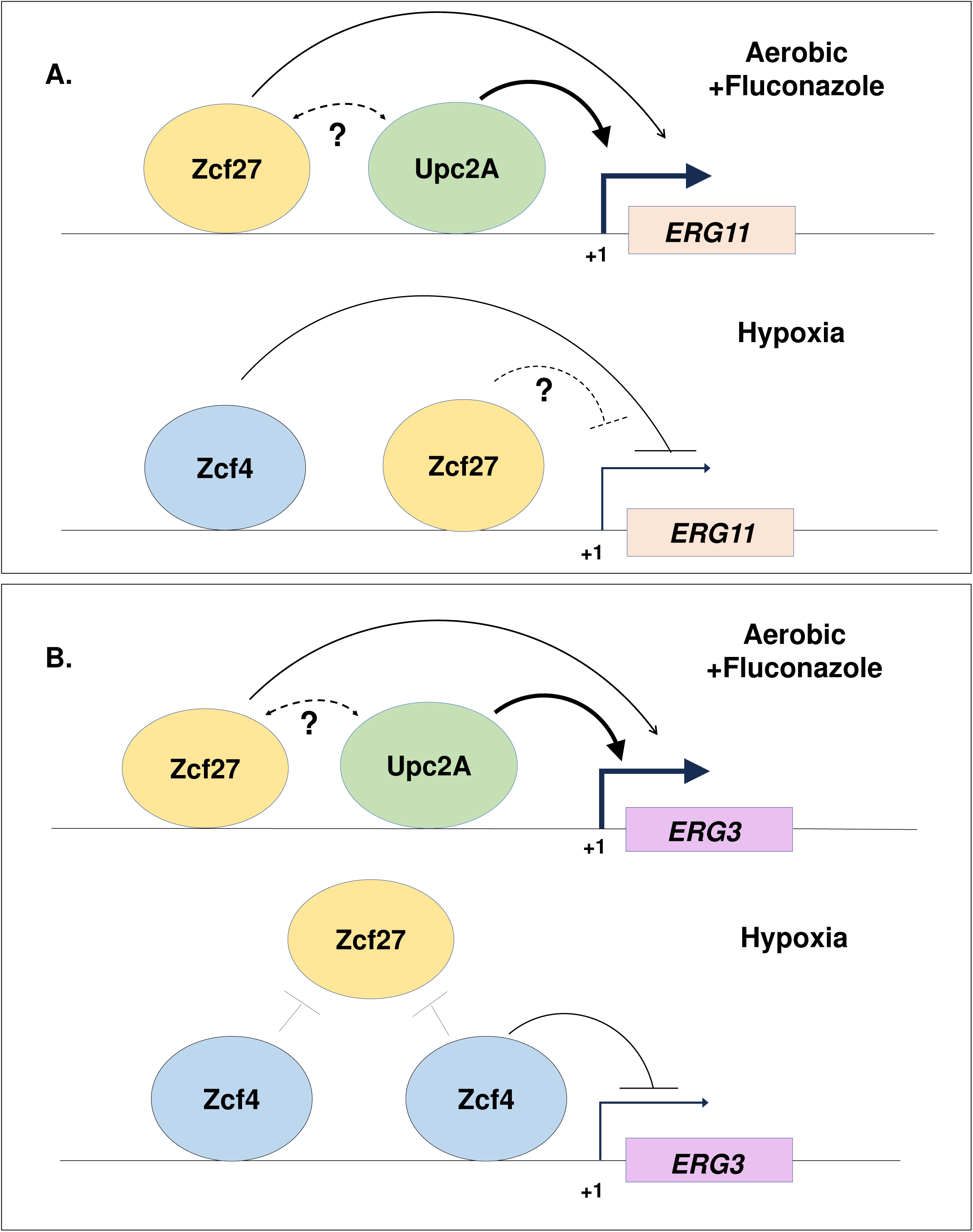
Model depicting the role of Zcf27 and Zcf4 in response to azole drug treatment and hypoxic conditions. **(A)** Under aerobic conditions with fluconazole treatment, Zcf27 (yellow) binds to the *ERG11* distal promoter region (E11P1), aiding in transcriptional activation of *ERG11*. Upc2A (green) binds to the *ERG11* proximal promoter region and is the commonly known transcription factor for *ERG11*. Zcf4 is not depicted or control expression of *ERG11* because it is not expressed, as determined by our data, under these conditions. Under hypoxic conditions, Zcf4 is induced and highly expressed where it binds to the distal promoter sequence of *ERG11* to repress *ERG11* and to prevent binding of Zcf27. Zcf27 binds to the proximal promoter likely to prevent Upc2A binding. **(B)** Under aerobic conditions with fluconazole treatment, Zcf27 (yellow) binds to the *ERG3* distal promoter region (E3P1), aiding Upc2A in transcriptional activation of *ERG3*. Upc2A (green) is known to bind to the *ERG3* proximal promoter region and a known transcription factor for *ERG3*. Again, Zcf4 is not shown to control the expression of *ERG3* because it is not expressed. Under hypoxic conditions, Zcf4 is induced and highly expressed where it binds to the distal promoter and proximal promoter regions of *ERG3* to repress *ERG3* and to prevent binding of Zcf27 and likely Upc2A. Overall, this indicates utilization of three zinc cluster transcription factors for direct and distinct promoter control of *ERG* genes in response to azole treatment and hypoxic conditions. For A and B, arrows represent activation, while bars indicate inhibition. The dotted lines and question marks denote unknown interactions or regulatory mechanisms.

Our phylogenetic analysis positions Zcf27 and Zcf4 as the closest homologs to *S. cerevisiae* Hap1 where we have determined that Zcf27 alters azole susceptibility, unlike Zcf4. Although Upc2A is the major transcription factor associated with azole-mediated induction of *ERG* genes, our study provides new insights into an additional transcriptional regulator besides Upc2 that is needed for azole-induced expression of *ERG* genes. Additional genetic and biochemical studies will be needed to determine the mechanism by which Zcf27 and Upc2A operate together in response to azole drugs. Nonetheless, we speculate that Zcf27 could mediate either a direct or indirect cooperative event that assists Upc2A in fully inducing ergosterol genes (see model Fig 12A and B). Additionally, in *S. cerevisiae*, deleting both Upc2 and its paralog Ecm22 further alters azole drug susceptibility, resistance to amphotericin B, and *ERG* gene expression (42, 43, 45, 65). Thus, Upc2A and Zcf27 may be operating in an analogous manner. However, there exists a distinct possibility that other yet-to-be identified zinc cluster transcription factors could be involved in regulating *ERG* gene expression. Identifying additional transcription factors besides Zcf27 and Upc2A will be important to fully understand what contributes to azole susceptibility and/or clinical drug resistance.

In contrast to Zcf27, Zcf4 protein levels were nearly undetectable under aerobic and/or azole treated conditions, with significant induction observed only under hypoxic conditions. This explains why the *ZCF4* deletion strain lacks an azole hypersensitive phenotype or any alteration in *ERG* gene expression. Based on our data, Zcf4 protein levels are likely being regulated by an unknown post-transcriptional mechanism. Although we have not identified the regulatory mechanism governing Zcf4 protein levels, we suspect that it is degraded via a specific ubiquitin ligase. Zcf4 may also be regulated in a manner similar to human HIF-1α (67, 68). To our knowledge, Zcf4 represents the first identified hypoxia-induced zinc cluster transcription factor and understanding the precise mechanism of protein degradation would be of interest.

Although deletion of Zcf4, also called Mar1 (Multiple Azole Resistance 1), has been initially described to alter azole susceptibility when treated with high concentrations of azoles, we have not been able to confirm this with our studies (34, 39). Currently, it is unclear the reason behind these discrepancies, but there could be differences in *C. glabrata* strains or conditions where Zcf4 is expressed at higher levels than what we have observed. However, the findings by Gale et al., utilizing a *C. glabrata* BG14 strain and employing a *Hermes* transposon approach to screen for fluconazole susceptibility, provided support for our observations (69). In their study, they identified several genes that when disrupted, altered azole drug susceptibility, including Zcf27 but not Zcf4 (69). More studies will be needed to completely understand the role of Zcf4 in azole susceptibility, if any, and how it is regulated at the transcriptional and post-transcriptional level. Nonetheless, our results are clear and consistent where Zcf4 plays a hypoxia-specific role in repressing *ERG* genes. We suspect that Zcf4, similar to Hap1 in *S. cerevisiae*, operates with a corepressor complex to repress *ERG* genes (59, 65, 66). Hypoxia-induced expression of Zcf4 and the growth defect observed in the *zcf4Δ* strain under hypoxic conditions highlight its importance in metabolic adaptation and survival in oxygen-limited environments. In addition, it is likely Zcf4 hypoxia-specific induction plays additional roles for *C. glabrata* to survive and propagate under low oxygen while within the humans.

Overall, this study expands our understanding of the transcriptional regulation of ergosterol biosynthesis in *C. glabrata*. This is significant because targeting ergosterol and/or enzymes involved in ergosterol biosynthesis have yielded highly useful and effective antifungals (66, 70, 71). Thus, studies focused on the regulatory mechanisms of this pathway could lead to the development of targeted antifungal therapies and help in overcoming the challenge of azole resistance in clinical settings. Because zinc cluster transcription factors are unique to fungi and not found in humans (32), there could be an opportunity to explore them as drug targets.

Overall, our findings reveal a novel regulatory mechanism where Zcf27 and Zcf4 are differentially employed by *C. glabrata* to manage ergosterol biosynthesis and maintain membrane integrity under varying environmental conditions. Our findings provide some of the first insights into functional role of two zinc cluster transcription factors. We suspect that further studies on these and similar factors will enhance our understanding of the pathophysiology and drug resistance mechanisms of *C. glabrata*.

## MATERIALS AND METHODS

### Plasmids and yeast strains

All plasmids and yeast strains are described in Table S3 and Table S4. The S288C BY4741 *S. cerevisiae* strain was obtained from Open Biosystems. The S288C strain containing the *HAP1-Ty1* sequence was corrected with a wild-type copy of *HAP1* (FY2609) and the *HAP1* deletion strain (FY2611) was kindly provided to us by Dr. Fred Winston, Department of Genetics, Harvard Medical School (40). *C. glabrata* 2001 (CBS138, ATCC 2001) and *C. glabrata* ATCC 200989 were acquired from the American Type Culture Collection (72). For Zcf27 complementation assays, a genomic fragment containing the *ZCF27* promoter, *5’ UTR*, open reading frame (ORF), and *3’ UTR* was PCR-amplified and cloned into the pGRB2.0 plasmid (Addgene) (73) using restriction enzymes BamHI and SacII. For endogenous C-terminal epitope tagging, a 3XFLAG-NatMX cassette was PCR-amplified from pYC46 plasmid (Addgene) and inserted at the C-terminus of *ZCF27* and *ZCF4* (74, 75). All *C. glabrata* strains were created using the CRISPR-Cas9 RNP system as previously described (74). Briefly, for generating deletion strains, two CRISPR gRNAs were designed near the 5′ and 3′ ORFs of the gene of interest. Drug-resistant selection markers were PCR-amplified using Ultramer DNA Oligos (IDT) from pAG32-HPHMX6 (hygromycin) or pAG25-NATMX6 (nourseothricin). For 3XFLAG epitope tagging, one CRISPR gRNA was designed in the *3′ UTR* of the gene of interest. Cells were then electroporated with the CRISPR-RNP mix and the drug resistance cassette.

### Serial-dilution spot and liquid growth assay

For serial-dilution spot assays, yeast strains were grown to saturation overnight in SC at 30°C. Cells were diluted to OD_600_ of 0.1 and allowed to grow to exponential phase with continuous shaking at 30°C. Each strain was then spotted in five-fold dilution starting at an O.D_600_ of 0.01 on untreated SC agar plates or plates containing 32 µg/ml fluconazole (Cayman). Plates were grown at 30°C for 2 days. For liquid growth assay, the yeast strains were inoculated in SC media and grown to saturation overnight. The cultures were then diluted to an OD_600_ of 0.1 and grown to log phase with shaking at 30°C. Upon reaching log phase, the strains were diluted to an OD_600_ of 0.001 in a 96 well round bottom plate containing 100 μL of SC media with and without 32 µg/ml fluconazole (Cayman). Cells were grown in liquid culture for 50 hours with shaking at 30°C, and the OD_600_ was measured every 15 minutes using a Bio-Tek Synergy 4 multimode plate reader. For spot assays under hypoxia, YPD plates were placed inside the BD Gaspak EZ anaerobe gas generating pouch system with indicator (BD 260683) after spotting and incubated for up to 7 days. Hypoxic cell collection for qRT-PCR, Western blot, and ChIP assays was performed by growing the indicated yeast strains in YPD media for 8 hours using the BD GasPak EZ anaerobe gas generating pouch system (BD 260683). Cells were immediately spun down for one minute and flash frozen to maintain the hypoxic state.

### Phylogenetic analysis

For the phylogenetic tree construction, 90 gene sequences were curated based on high-scoring BLAST hits to *Sc*Hap1. Of these sequences, 83 were retrieved from Mycocosm and 7 from the *Candida* Genome Database (76, 77), including CAGL0B03421g (*Cg*Zcf4), CAGL0K05841g (*Cg*Zcf27), B9J08_004061, B9J08_002924, B9J08_002930, B9J08_004353, and B9J08_002931. Protein sequence alignments were performed by Multiple Alignment using MAFFT version 7.471 (options: --auto) using the E-INSI iterative refinement method (78). The aligned sequences were then used to generate a maximum-likelihood phylogenetic tree with IQ-TREE version 1.5.5, using the built-in ModelFinder to determine the best-fit nucleic acid substitution model and 1000 ultrafast bootstrap replicates (79). The tree was visualized using Figtree software version 1.4.4 ( http://tree.bio.ed.ac.uk/software/figtree/).

### Quantitative real-time PCR analysis

RNA was isolated from strains grown in SC or YPD using standard acid phenol purification method. 1 µg RNA was reverse-transcribed to cDNA using the All-in-One 5x RT Mastermix kit (ABM). Gene expression primers were designed using Primer Express 3.0 software and are listed in Table S5. Quantitative real-time polymerase chain reaction (qRT-PCR) values are indicated in Table S7 and S9. At least 3 biological replicates, including three technical replicates, were performed for all samples. Data were analyzed by the comparative *C_T_* method (2*^−ΔΔCT^*) where *RDN18* (18S rRNA) was used as an internal control. All samples were normalized to untreated untagged wild-type strain. GraphPad Prism version 9.5.1 was used to determine the unpaired t-test for determining statistical significance.

### Yeast extraction and Western blot analysis

The indicated yeast strains were grown in SC or YPD media under aerobic or hypoxic condition. Yeast whole cell extraction and Western blot analysis to detect Zcf4-3XFLAG, Zcf27-3XFLAG and Histone H3 were performed as previously described (78). The monoclonal FLAG M2 mouse antibody (F1804, Sigma-Aldrich) was used at a 1:5000 dilution to detect Zcf4-3XFLAG and Zcf27-3XFLAG at 1:5000 dilution as previously described (59). The histone H3 rabbit polyclonal antibody (PRF&L) was used at a 1: 100,000 dilution as previously described (74).

### Chromatin Immunoprecipitation

Chromatin immunoprecipitation was performed using ZipChIP as previously described (79). Briefly, 50 mL cultures of indicated yeast strains were grown to exponential phase (OD_600_ of 0.6) in SC or YPD media with or without shaking at 30°C under aerobic or 8h of hypoxic condition, respectively. Cells grown in SC media under aerobic condition were treated with 64 µg/mL fluconazole (Cayman) for 3 h and collected. Cells were then formaldehyde cross-linked for 15 min and harvested as previously described (79). The cells were lysed by bead-beating with glass beads and lysate was separated from beads. Upon separation, cell lysates were transferred to Diagenode Bioruptor Pico microtubes and sonicated with a Diagenode Bioruptor Pico at the high frequency setting for 30 s ON and 30 s OFF for 20 cycles. After sonication, cell lysates were pre-cleared with 5 µl of unbound protein G magnetic beads (10004D, Invitrogen) for 30 min with rotation at 4°C. 300 µl of precleared lysate was immunoprecipitated with 10 µl of protein G-magnetic beads (10004D, Invitrogen) conjugated to 1 µl of M2 FLAG antibody (F1804, Sigma-Aldrich). Probe and primer sets used for qPCR analysis are described in Table S6, and qPCR values are indicated in Table S8 and S10

### Ergosterol extraction

Ergosterol was extracted from indicated strains as previously described (60, 80). Cultures were grown overnight in SC minimal media. Saturated cultures were back diluted to OD_600_ of 0.1 and were grown at 30°C to exponential phase (OD_600_ of 0.6), with or without 64 µg/ml fluconazole treatment. Sterols were extracted from yeast using 4 M potassium hydroxide in 70% (vol/vol) ethanol at 85°C for 1 h. After extraction, nonpolar lipids were separated by washing with methanol twice. Nonpolar sterols were crystallized after evaporating the n-hexane and dissolved in 100% methanol. Samples were analyzed by HPLC using a C18 column with a flow rate of 1 mL/min of 100% methanol. Ergosterol was detected at 280 nm, and cholesterol, used as an internal control for extraction, was detected at 210 nm.

## ACKNOWLEGEMENTS

This publication was supported by grants from the NIH National Institute of Allergy and Infectious Diseases to S.D.B. (AI136995) and to J.B.G (T32AI148103). Funding support was also provided from Purdue University AgSEED program (to S.D.B), Purdue Institute for Cancer Research (Grant P30CA023168: Bindley Metabolite Profiling Facility), Department of Biochemistry Bird Stair Fellowship (to D.S.), and NIFA 1007570 (to S.D.B).

